# Open spaced ridged hydrogel scaffolds containing TiSAMP surface chemistry promotes regeneration and recovery following spinal cord injury

**DOI:** 10.1101/2022.09.07.506969

**Authors:** Ahad M. Siddiqui, Fredric Thiele, Rachel Stewart, Simone Rangnick, Georgina Weiss, Bingkun K. Chen, Jodi Silvernail, Tammy Strickland, Jarred Nesbitt, Kelly Lim, Jean E. Schwarzbauer, Jeffrey Schwartz, Michael J. Yaszemski, Anthony J. Windebank, Nicolas N. Madigan

## Abstract

The spinal cord has poor ability to regenerate after injury, which may be due to cell loss, cyst formation, inflammation, and scarring. A promising approach to treat spinal cord injury (SCI) is the use of biomaterials. We have developed a novel hydrogel scaffold fabricated from oligo(poly(ethylene glycol) fumarate) (OPF) as a 0.08 mm thick sheet containing polymer ridges and a cell-attractive surface chemistry on the other side. When the cells are cultured on OPF with the chemical patterning, the cells attach, align, and deposit ECM along the direction of the pattern. Animals implanted with the rolled scaffold sheets had greater hindlimb recovery compared to the multichannel scaffold control, likely due to the greater number of axons growing across. Inflammation, scarring, and ECM deposits were equal across conditions. Overall, the results suggest that the scaffold sheets promote axon outgrowth that can be guided across the scaffold, thereby promoting hindlimb recovery.

## Introduction

The utilization of tissue engineering and material science in regenerative medicine for the treatment of spinal cord injury (SCI) presents an opportunity to modify the hostile microenvironment to promote regeneration after injury. Traumatic SCI occurs due to an insult of the vertebral column resulting in contusion, compression, laceration, or transection of the spinal cord. This primary injury leads to a host of secondary injury events, such as inflammation, cell loss, scarring, and cavitation, that not only creates a hostile microenvironment but a physical gap that impedes regeneration ^1^. Hydrogels can be constructed into scaffolding for axons to regenerate across, thereby bridging the injury. The macro and micro-architecture of the scaffold can greatly affect its ability to promote regeneration and recovery following SCI ^2^. Many considerations in the type of material or design play an important role balancing effects on regeneration verses inflammation or scarring because of the introduction of a foreign object.

Several scaffold macro-architectures designs have been investigated. These designs fall into 5 broad categories consisting of a spongey cylinder, single tube, channels, open-path with a core, and open-path without core ^3^. Open path scaffolds were shown to increase axon regeneration while maintaining the defect size, while other scaffold designs did not show as much axon growth and had increased defect size ^3^. However, other studies have demonstrated that multichannel designs can aid axon regeneration and have reduced cysts sizes, as well as can be designed to match the spatial distribution of spinal cord tracts ^4,5^. Another influence on axon regeneration is the diameter of the channels, where often reducing the channel size can increase axon count ^6^. For example, reducing the channel size from 660 μm to 450 μm in diameter nearly doubled the number of axons growing through ^6^. Channels as small as 200 μm have been shown to promote axon regeneration and *in vitro* characterization of astrocytes show that smaller diameter scaffolds alter alignment and proliferation ^7,8^. Another strategy to improve alignment uses grooves or ridges to introduce orientation through methods such as lithography ^9,10^, molding ^11,12^, or 3D biofabrication ^13,14^. Our group recently developed a method by which surface topology could be added to a water rich hydrogel through chemical processing utilizing hydrolysis ^15^. This surface chemistry uses a coating of titanium dioxide (TiO_2_) to layer a surface of phosphonic acid to create a self-assembled monolayer of phosphonates (SAMP; TiSAMP) that is only a few molecular layers high and can be patterned 30 μm wide. When patterned, fibroblasts extend and deposit ECM along the direction of the pattern ^16^.

One concern with implantable biomaterials, particularly as further modifications are made, is the biocompatibility of the material to the host tissue. It has been long observed that in matter of weeks, implant materials cause foreign body reaction that results in the accumulation of collagenous capsule and fibrosis with low vascularity and inflammation ^17–19^. One approach has been to reduce the area of the biomaterial available to interact with infiltrating cells or tissue, however this often leads to fibrosis at the tissue-biomaterial interface ^20,21^. Factors such as porosity, hydrophobicity, surface topography, stiffness, surface chemistry, and other active components can influence inflammation ^22^. For example, the ability of biomaterials to bind damage-associated molecular patterns (DAMPs) released from injured tissues may lead to greater inflammation around the biomaterial due to recognition by immune cells. As the material is degraded the DAMPs may re-release leading macrophage activation and chronic inflammation ^23,24^. Therefore, novel scaffold designs, modification, and biomaterials should be tested to ensure scarring or inflammation is not enhanced.

We have developed a hydrogel composed of oligo(poly(ethylene glycol)fumarate) (OPF) that is biodegradable, porous, and biomechanically similar to the spinal cord ^25,26^. When fabricated in a multichannel design it can reduce scarring and cyst formation and can induce greater axon outgrowth than poly(lactic-co-glycolic acid) (PLGA) and poly(caprolactone fumarate) (PCLF) based scaffolds ^26,27^. OPF can be easily utilized for combinatorial therapy by combining them with cells, drug eluting microspheres, extracellular matrix (ECM) proteins, or modifying the material with additional characteristics such as a positive charge or improved electroconductivity ^11,26–33^. All these modifications aim to improve regeneration and recovery following SCI. One important observation to come from these studies is that having directed axon growth across the scaffold is important and although the multichannel design helps achieve this, many of the axons still grow on the outside surface of the scaffold. In an aim to improve directed axon growth through the scaffold, we modified the design of the scaffold to introduce greater internal surface area by fabricating the scaffold as a ridged sheet ^11^. The ridges help to direct neurite growth in a bidirectional manner and when placed closer together can increase the number of neurites per surface area ^11^. This current study demonstrates that an open design, ridged scaffold sheet with a cell attractive TiSAMP chemical treatment (Supplemental Figure 1) improves axon outgrowth without increasing inflammation and scarring when compared to the more commonly used multichannel scaffold design. In addition, this study details a method by which machine learning algorithms can be used to analyze immune cell infiltration and scarring following scaffold implantation in a fast, non-bias manner.

## Methods

### OPF Sheet Fabrication

OPF was synthesized as previously described ^11,25,26^. Briefly, a liquid OPF polymer composed of 1g of OPF macromere (16,246 g/mol), 950μL deionized water, 0.3g N-vinyl pyrrolidinone (NVP; Sigma), and 0.05% (w/w) photoinitiator (Irgacure 2959; Ciba Specialty Chemicals) was made. A 0.08mm thin OPF sheet with 100μm high ridges was created using a Teflon mold with a 0.08mm spacer and grooves of 100μm depth spaced 1mm or 0.4mm apart. The polymer was slow pipetted onto the mold and covered with a glass plate. The mold was placed in a UV light oven for 1 hour and cured overnight in the dark. The single channel OPF scaffold was fabricated by mold injection of the polymer, cast over a 1mm wire. The single channel scaffold was then polymerized by exposure to an ultra-violet (UV) light (365nm) for one hour and cured overnight. The scaffolds were cut into 2mm lengths for transplantation. Similarly, for the 7 channel scaffold, the polymer was mold injected over 7 equally spaced 290μm wire in a glass cylinder and cut into 2mm lengths for transplantation.

### OPF Polymer Chemical Micropatterning

Chemical processing was done on the opposite side of the ridge surface of the OPF sheets (Supplemental figure 1) as previously described ^15^. Briefly, in a low atmosphere glovebox (OMNI-LAB Glove Box System, Vacuum Atmospheres Company) a solution of 30μl of titanium (IV) isopropoxide (CAS: 546-88-9) was mixed with 5ml anhydrous toluene (CAS: 108-88-3). A hydrolysis reaction created a surface of TiO_2_ on the hydrogel for bonding with phosphonic acid. The phosphonic acid solution is created by mixing 1mg of 12-(phosphonododecyl) phosphonic acid (CAS: 7450-59-1) with 5ml anhydrous toluene. OPF sheets are hydrated and positions on glass slides with the excess water blotted off. To create a pattern, a Kapton shadow mask with laser ablated slits of 30μm and spaced with 30μm material is placed on the OPF sheet. The sheet with the mask (or without the mask for fully coated sheets) is heated to 30°C for 15 seconds and placed in the TiO_2_ for 30 seconds followed by a rinse in toluene. The material is then heated to 30°C for 30 seconds. The material is then placed into the phosphonic acid solution overnight yielding a self-assembled monolayer of phosphonic acid (SAMP) on top of the TiO_2_ layer (TiSAMP). The scaffold is then rinsed in toluene, followed by isopropanol, and then distilled water. The mask is then removed and heated to 30°C for 30 seconds. The sheets are then kept in double distilled water until use.

### Human MSC isolation and culture

Human MSCs (hMSCs) were isolated from adipose tissue biopsies and then cultured, expanded, and cryopreserved by the Mayo Clinic Human Cellular Therapy Laboratory as previously described in ^34,35^. The release criteria included culture sterility, mycoplasma testing, cytogenetic analysis as well as cellular phenotyping including CD90+, CD105+. CD73+, HLA Class I+, CD14+, CD44+, HLA DR, and CD45-. The cells were cultured in advance MEM media (Gibco) supplemented with human platelet lysate (5%; Lawson, Mill Creek), 100 U/mL Antibiotic-Antimycotic (Gibco), 100 units/mL Glutamax (Gibco), and 1000 USP units heparin (Novaplus) at 5% CO2 and 37°C.

### Rat MSC isolation and culture

Rat MSCs were isolated from femur and tibia bone marrow from Sprague Dawley rats as previously described ^36^. They were cultured in in Dulbecco’s Modified Eagle Medium F12 medium (DMEM/F12; Gibco) supplemented with 10% fetal bovine serum, and 100 U/mL Antibiotic-Antimycotic (Gibco) at 5% CO2 and 37°C.

### Rat Schwann cell isolation and culture

Rat Schwann cells were harvested from the sciatic nerve of 2-5 day of Sprague Dawley pup as previously described 11.31.32,37,38. The cells were cultured in DMEM/F12 containing 10% fetal bovine serum, 100 U/mL Antibiotic-Antimycotic, 2 μM Forskolin (Sigma), 10 ng/mL neurgulinin-1 at 5% CO2 and 37°C.

### Co-culture hMSC and Schwann cells

For co-culture experiments, first the cells were grown alone as described above. Once 80% confluent they were culture on scaffold sheets at a 1:1 ratio (75,000 cells per cell type). The co-culture media of hMSCs and Schwann cells was composed of DMEM/F12 with L glutamine media (Gibco), Forskilin (2μM; Sigma), Neuroregulin (10 ng/ml; R&D), 10% FBS and 1% antibiotic/antimycotic (Gibco).

### Disassociated rat dorsal root ganglion neurons

Embryonic day 15 Sprague-Dawley rats were used to isolate dorsal root ganglions (DRGs; previously described in ^11,39,40^). Approximately 200 whole DRGs were pooled and then dissociated through mechanical trituration in trypsin. Each scaffold was plated with 50,000 cells in MEM supplemented with 15% calf bovine serum (Hyclone), 5 ng/mL nerve growth factor (Harlan Bioproducts), 7mg/mL glucose (Sigma Aldrich), and 1.2 mM L-glutamine (Gibco). To get a pure culture of neurons, the cells were treated with 1 x 10^-5^ M 5-fluoro-2-deoxy-uridine and with 1 x 10^-5^ M uridine (Sigma) for 3 days. Two days after treatment, the neurons were fixed in 4% paraformaldehyde in PBS for 30 minutes and washed 3 times in PBS before immunocytochemistry.

### OPF preparation and cell seeding

A positively charged OPF (OPF+) sheets were created from a polymer containing 1g of OPF macromere (16,246 g/mol), 650μL deionized water, 0.3g NVP, and 0.05% (w/w) photoinitiator, and 20% w/w 2-(methacryloyloxy)ethyl]-trimethylammonium chloride (MAETAC, Sigma-Aldrich). The positively charged OPF+ sheets were only used in the study of the cell interactions with topography and stiffness where chemical TiSAMP patterning was not introduced as a modification. The positive charge can help with cell attachment and outgrowth ^41^ but may interfere with chemical patterning. OPF or OPF+ was hydrated in culture media was pinned to sterile culture dishes containing SYLGARD 184 Silicone Elastomer (Dow Corning) to prevent spontaneous rolling of the scaffold. The plates were sterilized with 20-minute exposure to UV and multiple ethanol washes before pinning, while the scaffold sheets were sterilized using multiple isopropanol washes. The OPF sheets were coated with either media only, poly-l-lysine (PLL; Sigma-Aldrich; 0.01%; 5 minutes), laminin (100 μg/ml; Sigma-Aldrich), or rat plasma fibronectin (10 μg/ml) at 37°C overnight. Fibronectin was isolated from rat serum using gelatin-Sepharose affinity chromatography as previously described ^16^. Excess ECM protein coat or PLL was removed with 3 washes with culture media specific to the cell type that would be plated. Once the cells were 70-80% confluent, 150,000 cells were seeded onto each OFP sheet for 7 days. Co-cultured cells (50:50) were plated with 75,000 cells of each type. Three sheets per condition were used. The OPF sheets with the cells were fixed using 4% paraformaldehyde in phosphate buffered saline for 20 minutes. Three (3) washes with PBS were done to remove excess PFA and then immunocytochemistry was performed.

### Immunocytochemistry of cultured cells on scaffolds

Non-specific binding sites were blocked using a solution containing PBS, 10% normal donkey serum, and 0.3% Triton X-100 for 30 minutes. Then scaffolds containing cells were incubated with rabbit α fibronectin (1:100; R184), mouse α laminin (1:100; 2E8; Developmental Studies Hybridoma Bank ^42^), mouse α β-tubulin (1:300; Millipore) overnight at 4°C. Then the samples were washed 3 times with PBS and then incubated for 1 hour with the donkey α mouse Cy3 (1:200; Millipore Chemicon), donkey α rabbit Cy5 (1:200; Millipore Chemicon), and phalloidin conjugated to Alexa 488 (1:100; Molecular Probes) at room temperature. Then the samples were washed 3 times in PBS before being placed on a slide and coverslip in SlowFade Gold containing DAPI (Invitrogen). In the case of cells grown on coverslips, they were inverted onto a glass slide with SlowFade Gold containing DAPI following the same procedure above.

### Image acquisition for scaffold sheets

Fluorescent images were acquired using an inverted fluorescence microscope Zeiss Axio Observer Z-1 with a motorized stage mounted with an Axiocam 503 camera (Carl Zeiss). Zen 2 (blue edition; Carl Zeiss) software was used to acquire and process the images. Z-stacks with approximately 5 slides were taken at 20x. Bright field images of live cultures were taken on a Zeiss Axiovert Model 35 microscope with a Nikon CCD camera at 10x magnification. On each OPF sheet or coverslip, 4 representative areas were taken in a non-bias manner by moving one field of view up or down from a central position and 2 fields of view across. 3-4 different replicates were used for each condition (3-4 scaffolds or coverslips).

### Analysis of laminin and fibronectin deposit by Schwann cells and MSCs on scaffolds in culture

Fluorescent images were analyzed using ImageJ software (NIH). Fluorescence intensity was measured as previously described ^11^. Briefly, images type was set to 8-bit and the mean grey value of laminin and fibronectin was measured. The mean grey value of laminin and fibronectin was then normalized to the mean grey value of the DAPI staining per cell. A One-way ANOVA with Tukey’s multiple comparisons was performed to determine statistical significance. 4 representative areas were taken in a non-bias manner by moving one field of view up or down from a central position and 2 fields of view across. Four different replicates were used for each condition (4 scaffolds or coverslips).

### Scaffold preparation and surgical implantation

OPF sheets and single channel scaffolds were sterilized with serial dilution of isopropanol. Two (2) mm by 6 mm OPF sheets were rolled and placed into the single channel scaffold. Female Sprague Dawley rats (230 – 300 grams; Envigo) received a laminectomy at level T9 and a complete spinal cord transection was performed. The scaffold was placed in the transected area and all the muscles were sutured closed. The rats were divided into 4 groups that were implanted with either untreated rolled OPF sheets (OPF; n=7), fully coated TiSAMP OPF sheets (full TiSAMP; n=7), patterned TiSAMP sheets with 30 μm thick chemical and 30 μm thick polymer spacing (30×30 TiSAMP; n=7) or 7 channel OPF (n=8) scaffolds. The animals were cared for by veterinarians and other staff experienced with care of rats with spinal cord injury daily. All procedures were approved by the Mayo Clinic Institutional Animal Care and Use Committee and all guidelines were followed in accordance with the National Institute of Health, Institute for Laboratory Animal Research and the United States Public Health Services Policy on the Humane Care and Use of Laboratory Animals.

### Basso, Beattie, and Bresnahan (BBB) locomotor score

The BBB open field locomotor test was done weekly following injury to access hindlimb function. Two blinded observers rated aspects of the hind-limb function that was scored on a 21-point scale. The BBB scores were averaged between the observers as well as the left and right-side scores. Two-way ANOVA with Tukey’s multiple comparisons was used to determine statistical differences between the groups.

### Tissue preparation and Immunohistochemistry

The animals were euthanized by deep anesthesia and fixed by transcardial perfusion with 4% paraformaldehyde in PBS. The spinal column was removed en bloc and post fixed for 2 days at 4°C. The spinal cords were dissected out and a 1 cm segment with scaffold (T9) in the center was embedded in paraffin. Transverse sections 10 μm thick were made on a Reichert-Jung Biocut microtome (Leica, Bannockburn, IL).

The tissue sections were deparaffinized using serial dilutions of xylene and ethanol. Antigen retrieval was performed by immersing the slides in 1mM EDTA in PBS (pH 8.0) and then placing them in a streamer for 30 minutes. The tissue was then permeabilized with PBS containing 0.3% Triton X-100. Then blocking solution containing PBS with 0.3% Triton X-100 and 10% normal donkey serum was placed on the tissue for 30 minutes to block non-specific binding sites. Then the section were incubated in the primary antibodies, rabbit α fibronectin (1:100; R184), mouse α laminin (1:100; 2E8; Developmental Studies Hybridoma Bank ^42^), mouse α β-tubulin (1:300; Millipore), mouse α CD45 (1:50; DAKO), and rabbit α Iba-1 (1:800; Wako), overnight at 4°C. The sections were then serially washed with PBS containing 0.1% Triton X-100. The secondary antibody, donkey α mouse Cy3 (1:200; Jackson ImmunoResearch Laboratories) and donkey α rabbit Alexa 647 (1:200; Jackson ImmunoResearch Laboratories) was then placed on the sections for 1 hour at room temperature. The secondary antibody was washed off using PBS and then cover slipped with SlowFade Gold containing DAPI (Invitrogen).

### Trichrome Staining

Trichrome staining was done using a Masson Trichrome Staining Kit (Richard-Allan Scientific). The samples were first deparaffinized using serial dilutions of xylene and ethanol. The sections were then place in Bouin’s fluid for 1 hour at 56°C. After a 5-minute rinse in tap water, the slides were placed in working Weigert’s solution for 10 minutes. Following a 5-minute rinse in tap water, the slides were then placed in Biebrich Scarlet solution for 5 minutes. The slides were then rinsed in distilled water for 30 seconds followed by immersion into Phosphotungstic-Phosphomolybdic Acid for 5 minutes and Aniline Blue for 5 minutes. Then samples were then washed in 1% Acetic acid for 1 minute and rinsed in distilled water for 30 seconds. Lastly, the samples were then washed in 100% ethanol twice and xylene 3 times for I minute each. The slides were then mounted with mounting medium (Richard-Allan Scientific).

### Image acquisition

Immunofluorescent imaging was done using a Zeiss LSM 780 Confocal Microscope (Zeiss) with a motorized stage using Zeiss Zen blue edition software. Whole transverse sections were imaged using a Plan Apochromat 20x/0.8 M27 objective. Light microscopy was done using a Zeiss Axio Scanner Z1 (Zeiss) with the Zen Blue edition software. All sections on the slide were scanned using the automated tissue detector feature on the Zen software.

### Axon Count

Axons were identified by their characteristic punctate staining by β-III tubulin by a blinded investigator. Axons were counted on the whole transverse section at quarter lengths through the scaffold. The tiled 20x images were loaded into the QuPath Software^29^ and a grid was overlayed on the image. Using the point tool, the axons were manually counted by a blinded investigator. The axon was counted if it was in the quadrant or overlapped on the top horizontal or right vertical boundary. The area of the scaffold was done by tracing the scaffold boundary. The axons counts were then averaged through the quarter lengths of the scaffold and normalized to area for each rat. A one-way ANOVA with Tukey’s multiple comparisons was used to determine differences between the groups and the error was represented as ± SEM.

### Quantification of laminin and fibronectin components of the ECM in implanted scaffolds

A blinded investigator used ImageJ (NIH) to calculate the area of staining for fluorescent images with laminin and fibronectin staining through quarter length of the scaffold. First the region of interest around the scaffold was marked. The image was change to 8-bit and a threshold was set to select the staining in each channel. A percent area was calculated from the area stained to the area of the region of interest. The data was represented at quarter lengths or average through the scaffold regions for each rat. One-way ANOVA was done to determine differences between the conditions with the error represented as ± SEM.

### Machine learning analysis of immune cell infiltration and fibrotic scarring

For image analysis of the immune cell infiltration (CD45+ cells and Iba-1+ cells), a neural network was implemented in the QuPath software which represents a fast, unbiased and reproducible way to analyze microscopy images (Supplementary Figure 2). To identify the region of interest, a simple threshold pixel classifier was created to annotate tissue section. Next, we configured a watershed cell detection algorithm, to detect cells based on their intensity values of the DAPI channel. To enable a more accurate classification, additional features were calculated for each cell, including standard deviation of stain intensity for each stain, circularity, and perimeter. In the last step, we trained two separate networks on a large subset of training data, produced by manually annotating cells, for each antibody. Both Neural Networks were later combined into a composite classifier which could differentiate between CD45+ cells, Iba-1+ positive cells, cells positive for both markers, and cells negative for both markers. The data was then represented as an average of all the quarter lengths through the scaffold per rat or counts at each quarter length for each rat. A one-way or two-way ANOVA with Tukey’s multiple comparisons was used where appropriate.

For image analysis of the fibrotic collagen scar area, a color deconvolution based technique was implemented in QuPath for bright field images (Supplementary Figure 3). First the stain vectors were estimated using pixel values to separate out Analine blue stain from the Biebrich Scarlet Stain. A simple pixel threshold classifier was used (as explained for immunofluroscent images) to detect the tissue area. Then a color deconvolution was done to calculate the percent area of scarred tissue (more detailed methodology is available in the supplementary information). Measurements for whole tissue area and scar area were exported and percentages for fibrotic, scarred area were obtained by dividing scarred tissue area by whole tissue area. A one-way or two-way ANOVA with Tukey’s multiple comparisons was used where appropriate.

### Statistical Analysis

All data is reported as mean ± SEM and analysis was done using GraphPad Prism (GraphPad Software Inc.). Statistical significance was calculated using one-way or two-way ANOVA with Tukey’s multiple comparisons as appropriate. Graphical representation of the p value as represented as *p< 0.05, ** p< 0.01, *** p< 0.001, **** p< 0.0001, unless otherwise stated in the figure caption.

## Results

### Cultured mesenchymal stromal cells form spheres on ridged surfaces

OPF+ scaffold sheets were fabricated with 0.1 mm high ridges spaced 0.4 mm apart on one side. The other side contained bare polymer. Our previous study demonstrated that ECM protein coated ridged OPF+ sheets promoted Schwann cell and neuronal alignment and outgrowth along the ridge in vitro ^11^. However, different cell types respond differently to topographic cues (ridges or material stiffness), so 2 commonly investigated cell types for SCI treatment (mesenchymal stromal cells [MSCs] and Schwann cells) were compared. Human MSCs, rat MSCs, and rat Schwann cells were grown on the ridged OPF+ sheet and glass coverslips (Figure 1A-F). At 3 days in culture, the cells were found primarily on or near the ridges on the OPF+ sheet (Figure 1A-1C; white arrows indicate ridges). The Schwann cells adhered and spread along the ridges (Figure 1C), while human and rat MSCs did not spread but instead adhered as small or large cell clusters in the case of human MSCs (Figure 1A) or as individual cells (rat MSCs, Figure 1B). In contrast, when grown on glass coverslips (Figure 1D-1F), all cell types exhibit their normal, stereotypical, well-spread and flat shapes.

**Figure 1:**
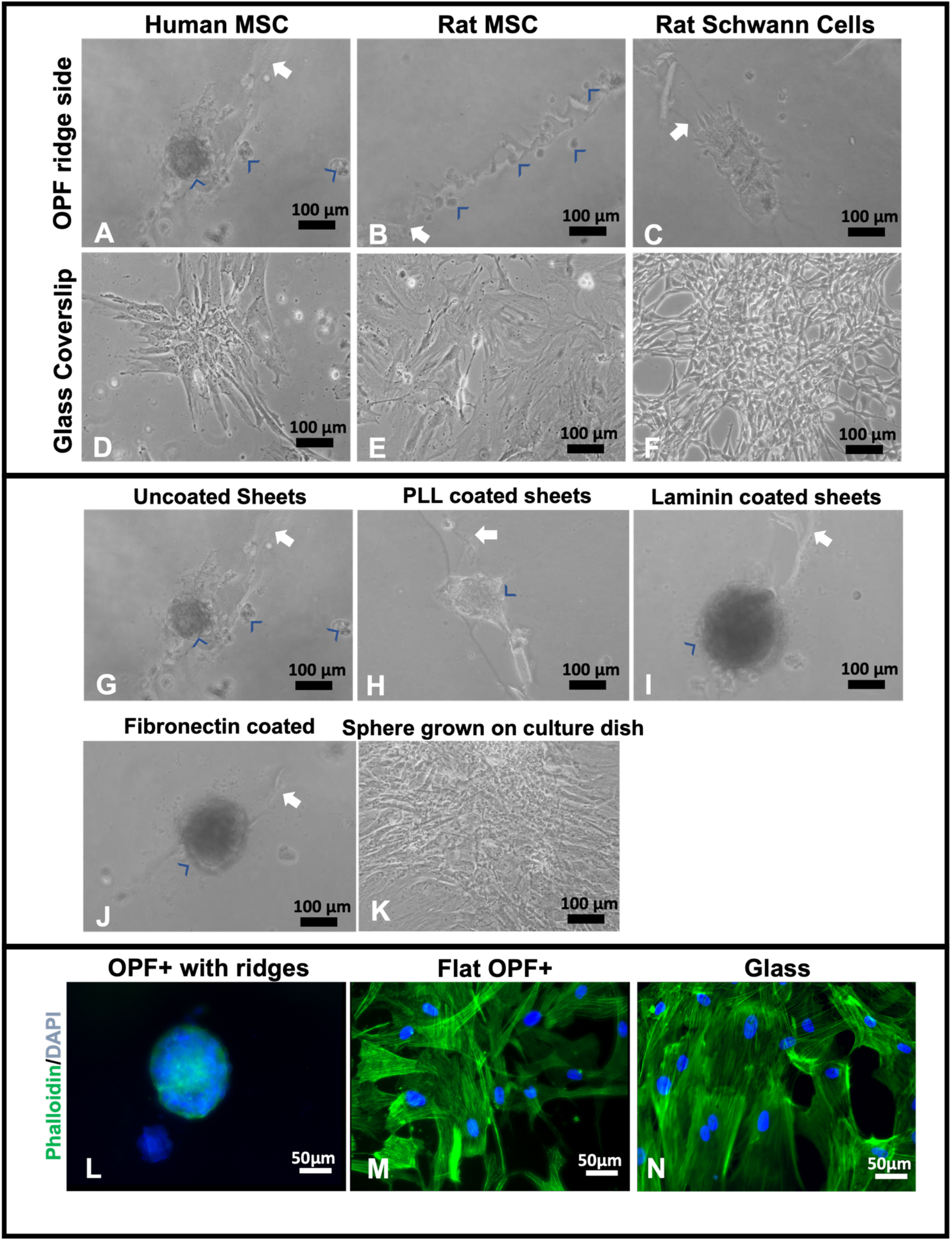
Box 1: hMSCs, rat MSCs, and rat Schwann cells grown on OPF+ scaffold sheets with ridges spaced 0.4 mm apart. Live culture images taken at 10x bright field. Human MSCs **(A)** and rat MSCs **(B)** attach but do not spread (blue arrowheads) when grown on ridged OPF+ sheets, whereas the rat Schwann cells attach and spread along the ridges (raised portions of the polymer are indicated by a white arrow) **(C)**. This is not observed on glass coverslips where hMSCs **(D)**, rat MSCs **(E)**, and Schwann cells **(F)** spread into their stereotypical morphologies. ***Box 2: hMSCs grown on coated OPF+ sheets containing ridges.*** Live culture images taken at 10x bright field. Human MSCs adhered in cell clusters (blue arrowheads) when grown for 3 days on uncoated **(G)**, poly-L-lysine coated **(H)**, laminin coated **(I)**, or fibronectin coated (J) OPF+ ridged sheets (white arrow indicates the ridge). If these spheres are picked and grown in a culture dish, they spread into their stereotypical shape **(K)**. ***Box 3: Staining for phalloidin and collagen IV of hMSCs cultured on OPF+ sheets with ridges, flat OPF+ sheets, or glass.*** hMSCs were grown for 7 days on OPF+ sheets containing ridges **(L)**, on the flat side of the OPF+ sheet **(M)** or on glass **(N)** and then fixed, permeabilized, and stained with DAPI (blue) to visualize cell nuclei, and phalloidin (green) for the actin cytoskeleton. On the ridged surface, multicellular clusters formed as indicated by multiple nuclei in a 3D spheroidal organization. On flat surfaces, cells attached and spread with similar morphology.

Since these OPF+ sheets were not coated with any ECM molecules other than those found in the media, we coated the OPF+ ridged sheets with PLL, laminin, or fibronectin protein coats. Again, we found that the hMSCs were found mostly on the ridges as cell clusters (Figure 1G-1J) 3 days in culture. To test if these clusters were viable and exhibit normal morphology, they were picked and plated in normal culture dishes (Figure 1K). The clusters were shown to spread on the culture dish and grow normally.

Next, we tested if this behavior of the cells was due to material stiffness or topography. When hMSCs are grown on the flat side of the OPF+ scaffold, we found that the cells adhered to the material in a flat manner like on glass and plastic (Figure 1M). Staining for actin with phalloidin, we show that the cytoskeletal structure looks like that of hMSCs grown on glass coverslips (Figure 1N). These results suggest that the 3D shape formed by the 0.1 mm high ridges induces cell clusters rather than the material being softer than plastic/glass.

### Mesenchymal stromal cells, Schwann cells, and DRG neurons attach to TiSAMP and extend along the chemical pattern

We fabricated ridged OPF scaffold sheets and deposited TiSAMP surface chemistry on the un-ridged side, either unpatterned (full TiSAMP) or patterned with 30 μm TiSAMP by 30 μm polymer (30×30 TiSAMP) using a shadow masking technique (Supplemental Figure 1). The chemical processing of the OPF hydrogel scaffolds with TiSAMP has the advantages of being only a few molecular layers thick and can be patterned closer together than molding ridges. The effect of the TiSAMP surface chemistry on rat MSCs, human MSCs, rat Schwann cells, neuronal outgrowth and ECM depositing was investigated. Human MSCs, Schwann cells, and dispersed DRG neurons were plated on OPF scaffolds with no TiSAMP, full TiSAMP coating, and patterned 30×30 TiSAMP for 7 days. The cells adhered in random orientation on the OPF (Figure 2A) and full TiSAMP (Figure 2B and 2D), although the cells adhered more sparsely on the untreated OPF. On the OPF 30×30 TiSAMP the cells aligned along the chemical pattern in a bidirectional fashion (Figure 2C). In some cases, the DRG neurons will extend on the bare polymer but then join at the contralateral pattern, still preferring the chemical’s bidirectional pattern as detected with anti-tubulin staining of neurites (Figure 2C and 2E). These results demonstrate that neurite outgrowth is enhanced on the TiSAMP material and alignment can be promoted through the striped pattern. Attachment and elongation are superior to that on the bare polymer and therefore may aid axon regeneration across the gap in the SCI animal model.

**Figure 2:**
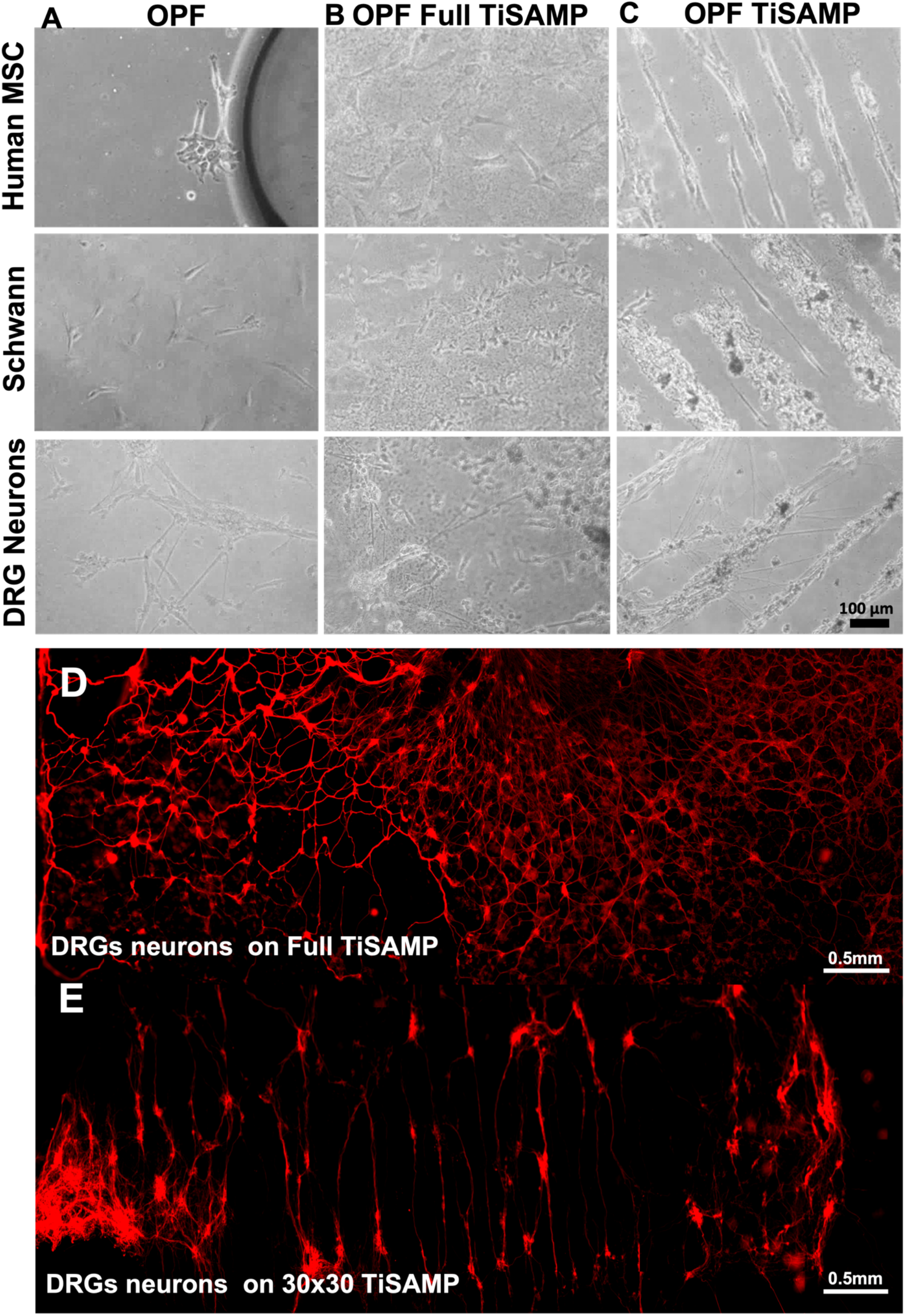
Attachment and alignment of hMSCs, Schwann cells, and dissociated DRG neurons on OPF scaffold sheets. Cells were plated on OPF scaffolds and grown for 5 days before staining and analysis. Cells on untreated OPF (A) showed no spatial directionality and were sparsely distributed 2 days in culture. (B) Cells on OPF scaffolds fully coated with TiSAMP (full TiSAMP) also demonstrated no spatial directionality, however, were more densely attached. (C) All 3 cell types aligned on chemically patterned OPF. DRG neurons sometimes branch out across the bare polymer to adjacent chemical strips that travel in a bidirectional orientation within 3 days of culture. Tiled micrograph of immunohistochemical staining for ß-tubulin of dispersed DRG neurons shows randomly distribution of neurons when plated on full TiSAMP (D) but highly organized on the patterned 30×30 TiSAMP scaffold (E) in a bidirectional orientation. Some axons are seen on the adjust polymer area but often find and orient on a different chemical strip.

The different cell types may interact with the biomaterial through depositing ECM. Human MSCs, Schwann cells, and a co-culture of hMSCs and Schwann cells were stained for fibronectin and laminin production. The co-cultured cells, like the cells cultured alone, aligned along the chemical pattern (Figure 3A-C). Human MSCs produce more fibronectin alone than Schwann cells, although not statistically significant (Figure 3D; hMSC = 17.02 ± 6.43 mean gray value and Schwann cells = 7.57 ± 2.87 mean gray value). Schwann cells produced more laminin (Figure 3E; hMSC = 43.33 ± 9.10 mean gray value and Schwann cells=161.9 ± 32.77 mean gray value, p=0.013). ECM deposits were in the orientation of the chemical patterning. Co-cultures can deposit both ECM types (Figure 3F). Co-culturing hMSCs and Schwann cells resulted in greater fibronectin deposit (188.9 ± 37.67 mean gray value) than Schwann cells (Figure 3J; p=0.0004) or MSCs alone (p=0.0003) suggesting that both cell types are producing fibronectin. Co-culture staining for laminin was not significantly different from Schwann cells alone but was much higher than hMSCs alone (Figure 3K; hMSCs = 43.33 ± 9.10 mean gray value, co-culture = 147.50 ± 17.97 mean gray value, p=0.0367) indicating that Schwann cells are primarily responsible for producing laminin in this system. DRG neurons were plated onto Schwann cell cultures and DRG neurites were visualized by anti ß-tubulin staining which shows some neurites extended along the fibronectin component of the ECM (Figure 3L). Together, these results indicate this TiSAMP-patterned material is able to support cell attachment, neurite outgrowth, and ECM deposition, all of which are important for SCI repair.

**Figure 3:**
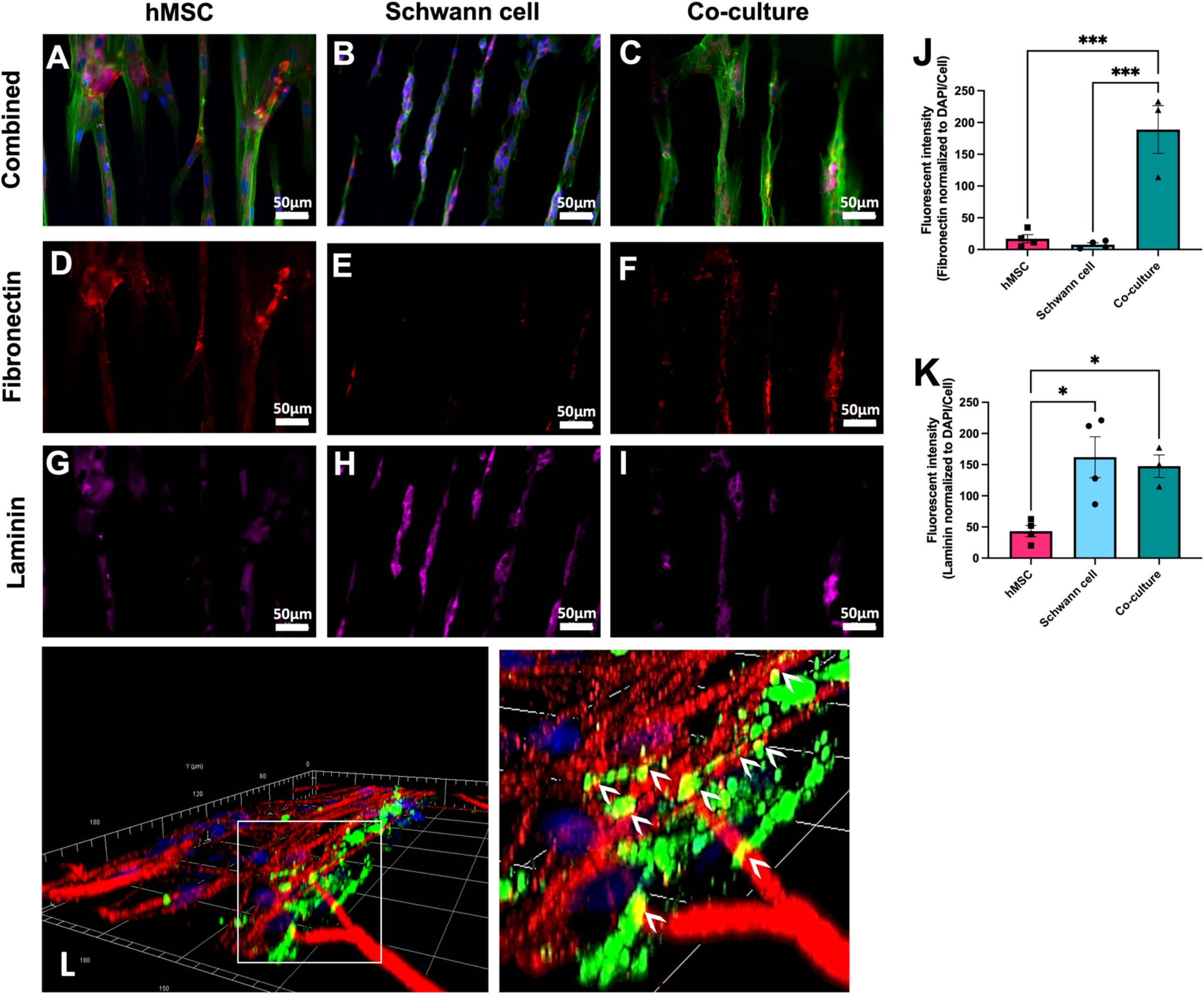
Attachment, polarization, and ECM deposit by hMSCs and rat Schwann cells on patterned 30×30 TiSAMP OPF. hMSCs (A) and Schwann cells (B) were cultured on the chemically patterned OPF individually and in co-culture (50:50) on the chemically patterned OPF for 7 days. Using immunocytochemistry, the scaffolds were stained for DAPI (blue), Phalloidin (green; A-C), Fibronectin (red, D-F) and Laminin (pink, G-I) to compare ECM production. Fibronectin (J) and laminin (K) intensity was quantified using image J by normalizing Fibronectin production to DAPI/cell number (n=4 scaffolds with 4 areas per sheet). Statistical analysis of Fibronectin staining was run using a one-way ANOVA with multiple comparisons using Tukey’s post-hoc test. The degrees of significance are indicated as follows: p<0.05 (*), p<0.01 (**), and p<0.001 (***). Error bars represent standard error. (L) A 3D reconstruction of a confocal z-stack image is shown containing a grid with each square representing 30×30 microns (L left) and a zoomed-in section (L right). Samples were stained for fibronectin (green), β-tubulin (neurons, red) and DAPI (cell nuclei, blue) after Schwann cells and neurons had been in culture for 10 and 4 days, respectively. Neurites were observed to weave over and through (white arrowheads) the network of the native, SC-specific ECM.

### Rolled scaffolds have equal number of hemopoietic cells, microglia, and other cell types as multichannel scaffolds

The in vitro data demonstrates that TiSAMP is cell and protein (ECM) attractive as more can bind to the surface than on bare OPF surfaces. While enhancement of axon regeneration is an important goal for treatment of SCI, any modifications (chemical or architectural) to implanted polymer scaffolds should not lead to increased negative side effects. Foreign body reactions leading to inflammation and scarring have been a long concern for neuroprosthetics and biomaterials ^31,32,43,44^. The cell and protein attractive surface may lead to greater cell infiltration into the scaffold resulting in enhanced inflammation or fibrosis/scarring. Immunostaining was done with CD45 to label hemopoietic cells, Iba1 to label with microglia/macrophages, and DAPI counterstain to label all cell nuclei to determine the infiltration of cells into the scaffold after transplantation (Figure 4A-D). A machine learning approach was used to identify and quantify the cell populations in a quick and non-biased manner (details in Supplementary Figure 2). All animals implanted with OPF scaffolds had equal numbers of hemopoietic cells, microglia/macrophages, and total number of cells equally distributed through the lengths of the scaffolds. The average number of hemopoietic cells was 82.88 ± 21.37 cells/mm^2^ for untreated OPF, 50.12 ± 5.19 cells/mm^2^ for full TiSAMP, 79.11 ± 19.90 cells/mm^2^ for 30×30 TiSAMP, and 91.65 ± 33.74 cells/mm^2^ for the 7 channel scaffolds (Figure 5E). Microglia/macrophage were 73.78 ± 27.13 cells/mm^2^ for untreated OPF, 55.66 ± 23.96 cells/mm^2^ for full TiSAMP, 90.57 ± 27.25 cells/mm^2^ for 30×30 TiSAMP, and 120.4 ± 37.34cells/mm^2^ for the 7 channel scaffolds (Figure 4F). Interestingly, these immune cell types only consisted of a small proportion of the total cells which averaged 4592.4 ± 221.0 cells/mm^2^ for untreated OPF, 3956.3 ± 166.4 cells/mm^2^ for full TiSAMP, 3558.6 ± 375.4 cells/mm^2^ for 30×30 TiSAMP, and 4588.5 ± 80.9 cells/mm^2^ for the 7 channel scaffolds (Figure 4G). These results show no statistical differences in the types and numbers of cells that interact with these different scaffolds after implant.

**Figure 4:**
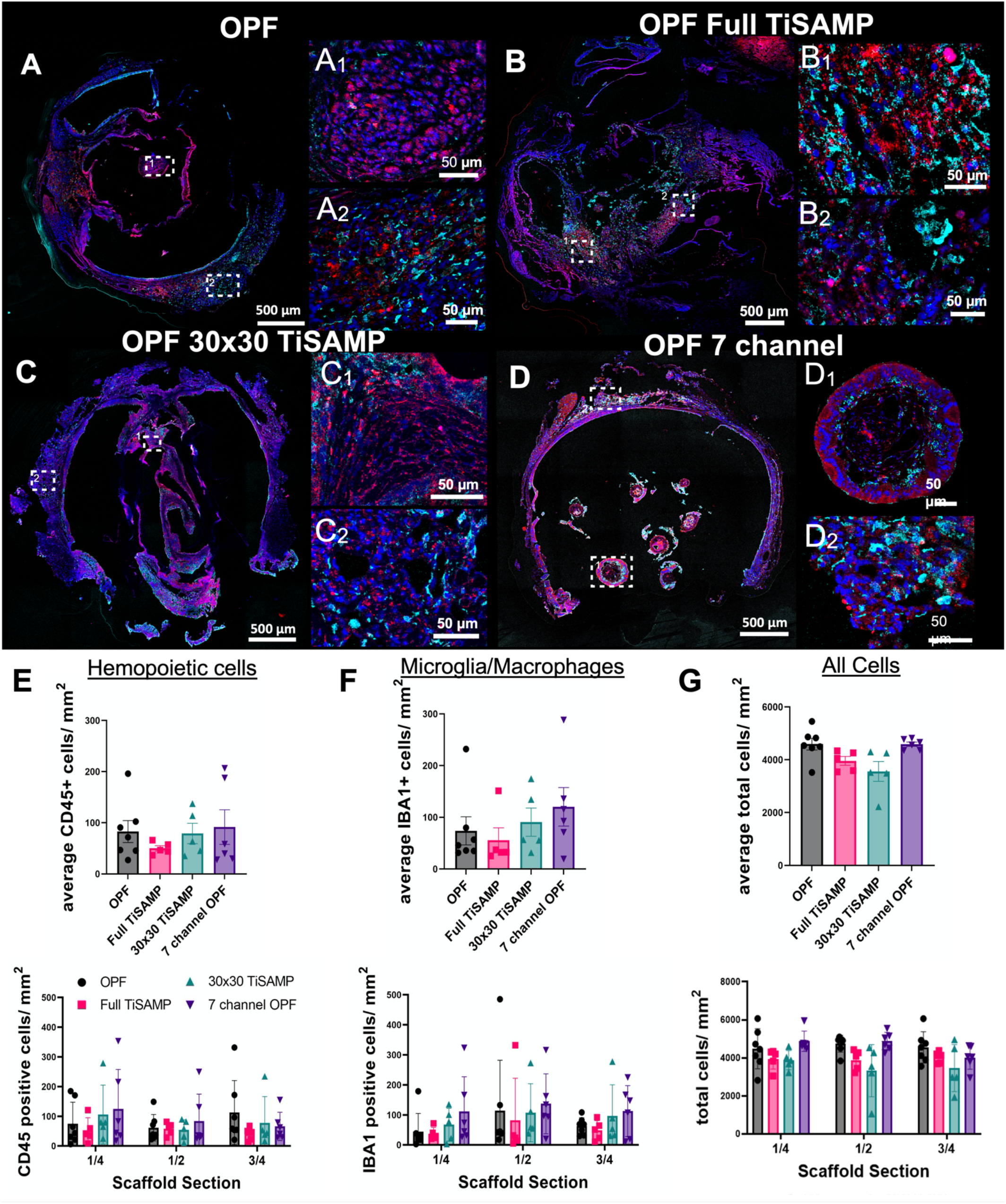
Infiltration of immune cells into the implant OPF scaffolds. Implanted untreated OPF (A), Full TiSAMP (B), 30×30 TiSAMP (C), and 7 channel (D) scaffold were sectioned and stained at quarter lengths (¼, ½, and ¾ of the scaffold; center or ½ segment shown here) for Iba-1 (red), CD45 (cyan), and DAPI (blue) to determine infiltration of microglia (E), hemopoietic cells (F), and all cells (G) respectively. Magnified segments of the tiled image are shown for a central portion of the scaffold (A1-D1) and the outer portion (A2-D2). A machine learning algorithm was designed to determine the number of microglia (E), hemopoietic cells (F), and all cells (G) as well as the total area of the scaffold. There was no significant difference between the conditions. One-Way ANOVA with Tukey’s multiple comparisons.

**Figure 5:**
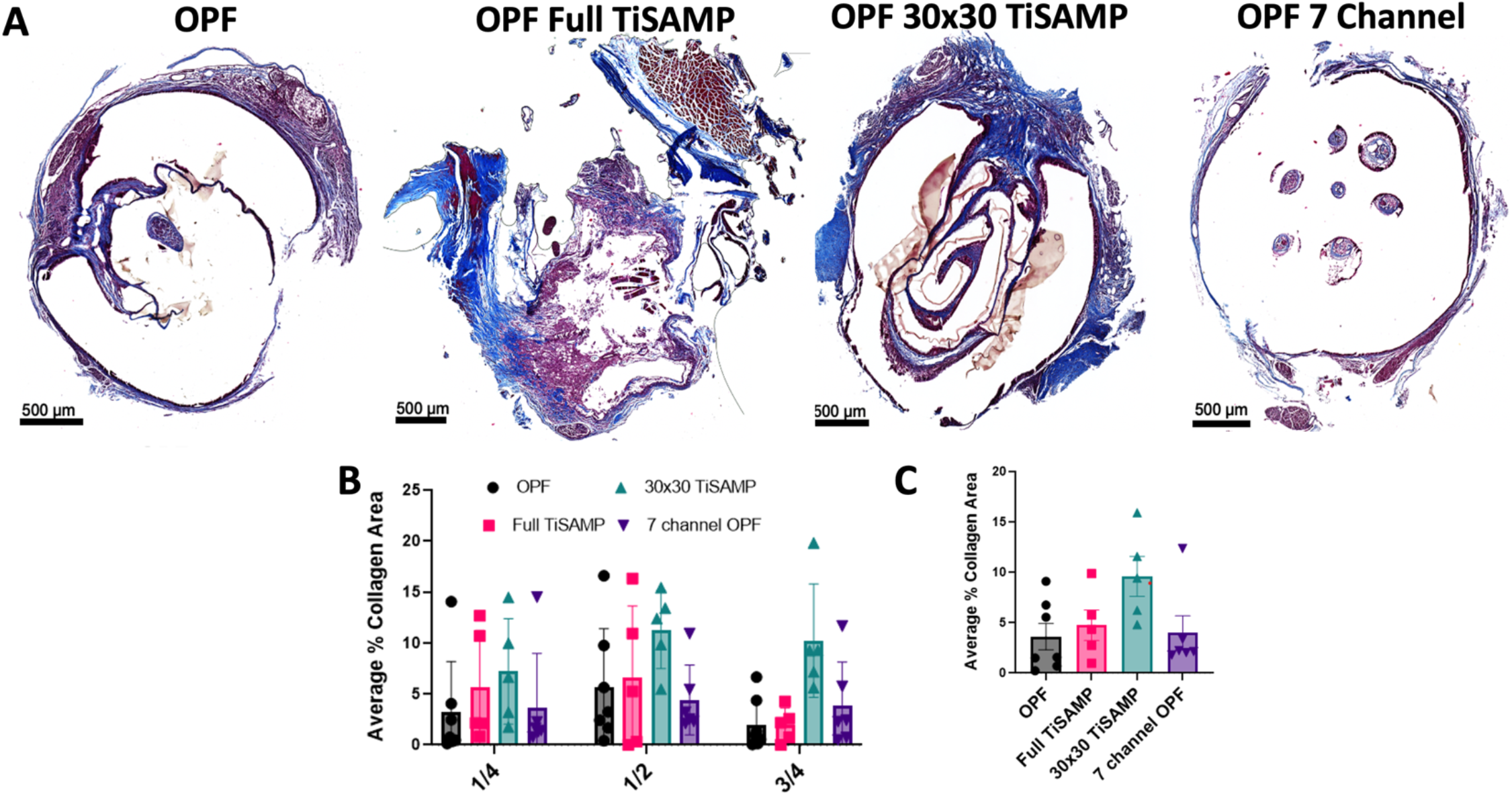
Fibrotic scarring following implantation of OPF scaffolds. (A) Implanted untreated OPF, Full TiSAMP, 30×30 TiSAMP, and 7 channel scaffolds were sectioned and stained at quarter lengths (¼, ½, and ¾ of the scaffold; center or ½ segment shown here) with Trichrome histological stain. The collagen component is indicated by blue staining whereas the cytoplasm is pink. The dark blue staining represents areas containing excessive collagen deposits characteristic of fibrotic scarring with the light purple being an overlap of regular collagen deposits within the tissue. A machine learning algorithm was used to analyze the average percent area of collagen scarring at quarter lengths of the scaffold (B) and then averaged (C). There was no significant difference between the conditions. One-Way ANOVA with Tukey’s multiple comparisons.

### Rolled scaffolds have equivalent deposits of collagen, laminin and fibronectin as multichannel scaffolds

The extra open space could have been recognized as cavitation or the chemical processing could have been detected as an additional foreign object resulting in greater scarring. In addition, it has been noted that cell infiltrate into the scaffold and may contribute by laying down ECM on the TiSAMP surfaces. Trichrome staining was done to determine the amount of collagen fibrotic scarring present in the rolled scaffolds compared to the 7 channel scaffolds (Figure 5A). A machine learning approach was employed to recognize whole tissue and the scarred area from whole slide scans (details in Supplementary Figure 3). There was no statistical difference in the amount of collagen scar present across the scaffold types with a roughly equal distribution across the length of the scaffold (Figure 5B and C). The average percentage of scar area present was 3.62 ± 1.31% for untreated OPF, 4.73 ± 1.51% for full TiSAMP, 9.58 ± 1.98% for 30×30 TiSAMP, and 3.79 ± 1.69% for the 7 channel scaffolds (Figure 5C). Large amount of the scar was observed on the outer surface of the rolled scaffolds rather than internally.

Since cultured cells can deposit ECM on the chemical pattern which neurites can grow on and weave through (see Figure 3), the infiltrating cells may also contribute to the amount of ECM on the chemically processed scaffolds. Fibronectin (Figure 6 A, D, G, J) and laminin (Figure 6 B, E, H, and K) are some of the major ECM types in the spinal cord. Equivalent amounts of fibronectin and laminin were detected across the lengths of the scaffolds independently of whether they were chemically treated or untreated scaffolds. The total contribution of ECM to the surface areas of the scaffolds was 10-20%. This indicates that the presence of the TiSAMP coat did not enhance or inhibit ECM depositing.

**Figure 6:**
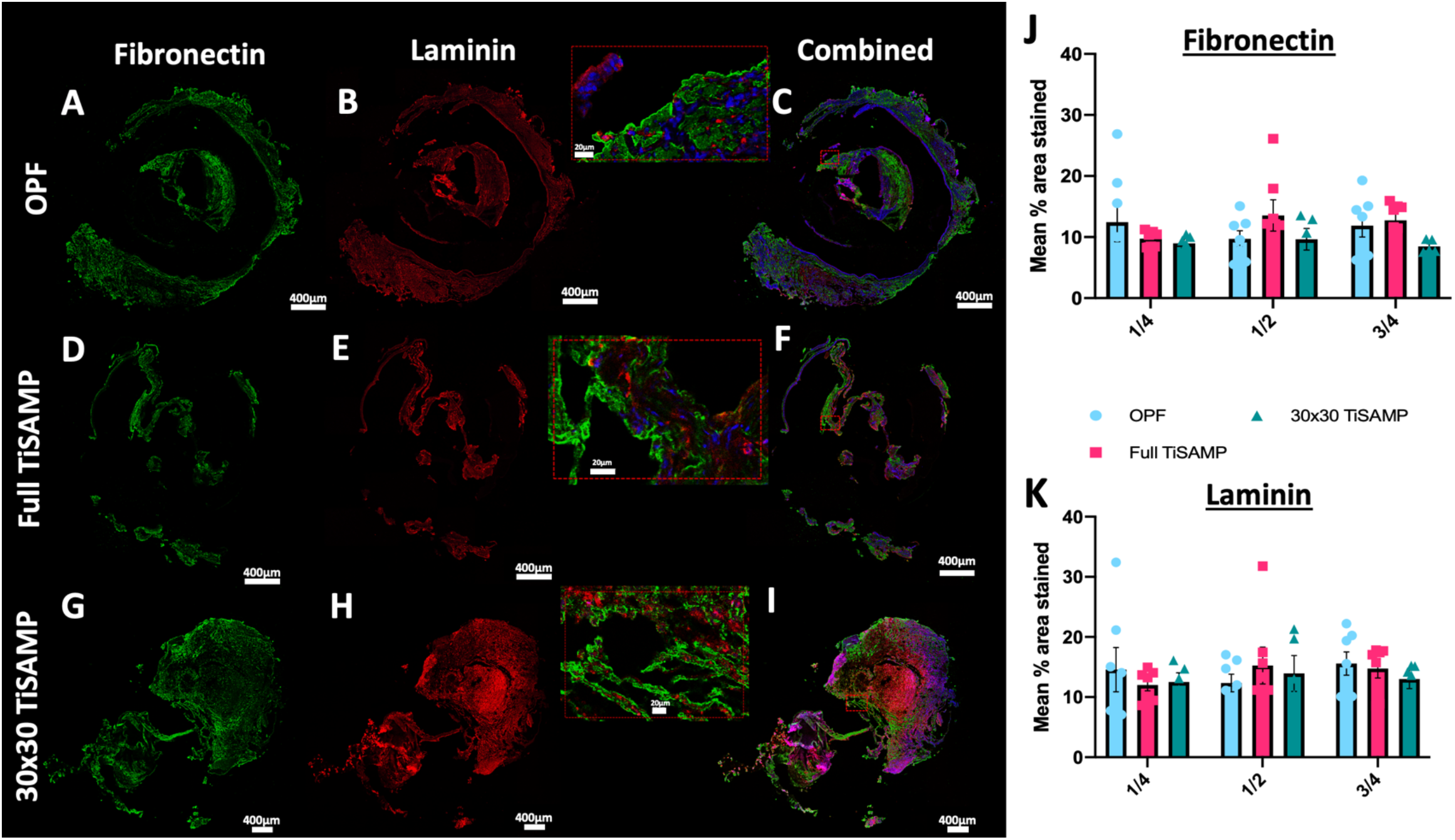
Fibronectin and Laminin deposits by infiltrating cells following implantation of rolled OPF scaffolds. Quarter lengths (¼, ½, and ¾ of the scaffold; center or ½ segment shown here) of untreated rolled OPF scaffold (A-C), full TiSAMP OPF (D-F), and 30×30 TiSAMP (G-I) were stained for fibronectin and laminin. Magnified segment of the region of interest in the combined images (C, F, I) are shown in the red box. There was no significant difference in the mean percent area stained through the whole length of the scaffold (One-way ANOVA with Tukey’s multiple comparisons) of either fibronectin (J) or laminin (K).

### Implantation of rolled scaffolds results in greater axon growth through the scaffold and improved hindlimb function compared to multichannel scaffolds

Chemically treated ridged OPF sheets (0.4mm spaced ridges on one side and either no chemical treatment, full TiSAMP treatment, patterned 30×30 TiSAMP treatment on the other side) were rolled into a single channel OPF scaffold and implanted into rats following thoracic (T9) transection. The effect on axon outgrowth, inflammation, scarring, and behavioral recovery was compared between animals implanted with rolled OPF sheets and OPF in a multichannel (7 channel) design in the absence of any added ECM or Matrigel over a 5-week period. Immunostaining of axons with ß-tubulin (Figure 7 A-D) revealed that all 3 rolled scaffolds (untreated OPF contained 2936 ± 580.2 axons/mm^2^, p=0.0091; Full TiSAMP had 2579 ± 611.0 axons/mm^2^, p=0.0243; 30×30 TiSAMP had 1733 ± 949.0 axons/mm^2^, p=ns) had greater axon counts than the 7-channel scaffold (125.9 ± 26.92 axons/mm^2^, Figure 7E). Much of the ß-tubulin staining in the 7-channel scaffold were cells that contained ß-tubulin, possibly though digestion of axonal/neuronal debris (axons are identified by punctate ß-tubulin staining with no DAPI nuclei whereas cells are indicated by ß-tubulin staining containing DAPI nuclei; Figure 7D). Although there is an increase in the number of axons in the scaffold, axons growing on the outside surface were still evident (Figure 7 A1-C1). The rolled sheet design, with more open surface area available in the same size scaffold as the 7-channel design, contained more axons per mm^2^ perhaps indicating a greater regenerative potential to make connections across the lesion.

**Figure 7:**
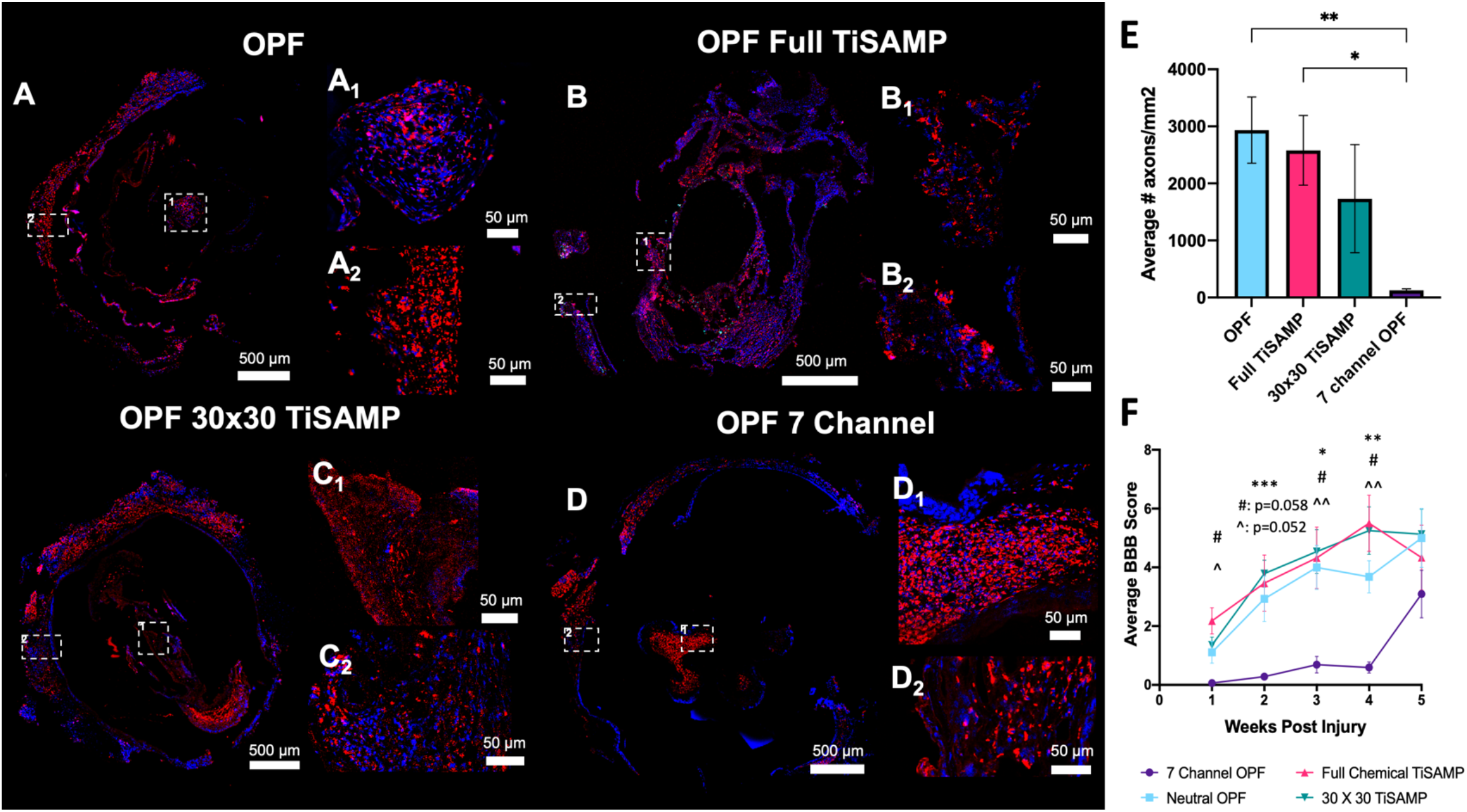
Axon outgrowth into rolled scaffold with or without TiSAMP compared to 7 channel scaffolds following transection and implantation into the T9 spinal cord. Implanted untreated OPF (A), Full TiSAMP (B), 30×30 TiSAMP (C), and 7 channel (D) scaffold were sectioned and stained at quarter lengths (¼, ½, and ¾ of the scaffold; center or ½ segment shown here) for β-tubulin (red) and DAPI (blue) to determine axon growth. Axons were determined by punctate staining positive for β-tubulin whereas cells were β-tubulin positive with DAPI nuclei (likely phagocytosing cells) and were not counted as axons. Magnified segments of the tiled image are shown for a central portion of the scaffold (A1-D1) and the outer portion (A2-D2). The average number of axons growing through the rolled scaffold was greater than the 7 channel scaffolds (E; One Way ANOVA with Tukey’s multiple comparisons; error bars = SEM; degrees of significance are indicated as follows: p<0.05 (*), p<0.01 (**)). The average hindlimb function as measured by the BBB motor test also indicated that there was early and sustained hindlimb functional improvement when the rolled scaffold was implanted compared to the 7 channel scaffold (purple line; Two-Way ANOVA with Tukey’s multiple comparisons; degrees of significance are indicated as follows: p<0.05 (*), p<0.01 (**), p<0.001 (***); * untreated rolled OPF compared to 7 channel OPF, # 30×30 TiSAMP OPF compared to 7 channel OPF, ^ full TiSAMP OPF compared to 7 channel OPF).

The BBB test is an open field test that measures hindlimb function based on a 21-point score with 21 being normal function, score of 8 being able sweep with no weight support, and 0 being no function. The BBB score was averaged from 2 blinded scorers weekly over the 5-week period. The BBB scores matched the trend from the axons counts where animals implanted with either of the 3 rolled scaffolds had significantly greater scores from week 1 to 4 than those animals implanted with the 7-channel scaffold (Figure 7F). The greatest difference in the BBB score was seen at week 3 (7 channel scaffold score was 0.69 ± 0.28 vs. untreated OPF score of 4.00 ± 0.74, p=0.015; Full TiSAMP score of 4.32 ± 1.05, p=0.048; 30×30 TiSAMP score of 4.54 ± 0.75, p=0.006) and week 4 (7 channel scaffold score was 0.59 ± 0.19 vs. untreated OPF score of 3.68 ± 0.55, p=0.004; Full TiSAMP score of 5.50 ± 0.96, p=0.012; 30×30 TiSAMP score of 5.25 ± 0.81, p=0.007). The animals implanted with the 7 channel scaffolds improved at week 5 to the point that the greater BBB scores of the rolled scaffolds were not significantly greater. These results suggest that a rolled scaffold design may facilitate more rapid recovery of function perhaps by increasing the number of axons able to traverse the scaffold.

## Discussion

Anatomical and functional recovery following SCI requires a growth permissive environment that guides axons across the injury to help bridge the gap and reform connections. Both short relay reconnections and regeneration of long tracts can promote functional recovery ^45^. One way to promote these reconnections is to provide appropriate growth environments with topographical cues, which can be achieve with the introduction of hydrogel scaffolds. This study demonstrates that chemically patterned TiSAMP OPF sheets can promote alignment and outgrowth of cultured hMSCs, Schwann cells, and dispersed DRG neurons. The ECM matrix produced by these cells can also align on the patterned surface and the use of multiple cell types can contribute different ECM types. *In vivo*, rats implanted with the open space rolled scaffold sheets with ridges had greater axon regrowth through the scaffold with improved early hindlimb recovery compared to rats implanted with the multichannel scaffold without any increased inflammation, scarring, or ECM (laminin and fibronectin) deposition. Lastly, research in this space is slowed down by the in-depth histological analysis often required. We demonstrated a method by which machine learning utilizing neural networks can aid in analysis of these data sets in a fast, non-bias manner allowing for rapid characterization of implanted materials using an open-source software, QuPath.

Notably axon regeneration, even without the aid of ECM molecules, cells, or other factors, was greater in the rolled scaffold sheet design than the multichannel design. Previous studies have demonstrated that open path polycaprolactone (PCL) scaffolds increased axon regrowth and had smaller deficit size when compared to multichannel, tube, or cylinder designs ^3^. However, mutlichannel scaffolds with increased number of smaller channels can encourage greater axon regrowth as well ^6^. This could partially be due to the larger channel scaffolds having larger fibrous rims. Our rolled scaffold design has increase surface area to that of the multichannel design and is somewhat a hybrid of an open path and multichannel scaffold since the ridges do act as spacers between the layers and segment the open space. The rolled scaffolds with its greater open spaces did not result in any greater amount of fibrotic collagen scarring than the multichannel scaffolds. Additional topographical cues are added with the TiSAMP chemically patterning that in vitro helps to align and guide growth of different cell types. This approach is most like the use of fibrils or grooves micro-architectures to promote elongation and alignment ^11,46^. A PLGA scaffold designed as a cylindrical nanofibrous scaffold was demonstrated to help align cells and induce their proliferation in vitro and after transplantation in a left lateral hemisection had superior hindlimb function to lesion control animals on the BBB open field test ^47^. Our study demonstrated similar level of hindlimb function recovery when the rolled OPF scaffold sheet was implanted following complete transection. Although, in the absence of added factors such as ECM or cells, all 3 rolled scaffold types had similar level of recovery and axon regrowth through the scaffold. However, all of the rolled scaffold sheets in this study have ridges that we have previously shown can guide neurites in vitro alone ^11^. Future studies should explore if adding aligned ECM or cells, which is aided by the TiSAMP patterning, could further enhance regeneration and recovery.

MSCs are a popular cell type for use in CNS injuries and disease. They can provide trophic support, modulate the immune system, aid tissue preservation, and promote angiogenesis ^48,49^. On ridged surfaces, MSCs preferentially adhere to the ridges, instead of the adjacent flat area, however forming spheres. On flat OPF sheets, glass, or plastic, MSCs will take their stereotypical flat morphology. The addition of ECM protein coats did not aid in flat attachment to the ridged OPF scaffold. The MSCs produced collagen IV on the softer, spinal cord like OPF material. When MSCs are grown on the OPF surface containing patterned TiSAMP that was only a few molecular layers higher than the polymer spacing, they attach and align on the scaffold without forming spheres. Taken together, this demonstrates the importance of the stiffness and physical structure of surfaces where the MSCs are expected to adhere or engraft. When used for treatments for CNS injures and disorders, cells are injected into or on top of soft surfaces such as the brain or spinal cord. However, many of the studies that are done to characterize MSCs *in vitro* are done on glass or plastic surfaces ^50–52^. In this study we demonstrate that cell characteristics can change depending on surface morphology and stiffness. Gene expression changes, cell adhesion properties, and changes in proliferation have been previously described being influenced by surface structure and stiffness that the MSCs are grown on top ^53,54^. This scaffold with or without TiSAMP introduces a new way to investigate cells on surfaces closer to the biomechanics that they will be implanted in. In addition, the scaffold also presents a way to deliver pre-cultured cells and ECM to the spinal cord in future studies, while maximizing surface area and promoting bidirectional orientation to bridge the gap after injury.

Schwann cells are another popular cell therapy choice for SCI as they are the myelinating cell type of the peripheral nervous system and may encourage regeneration through secretion of trophic factors, ECM proteins, and physical scaffolding ^1,55,56^. However, the amount of ECM produced by Schwann cells may be limited and in the current study produced little fibronectin. Other cell types, such as fibroblasts have been used to enhance depositing of factors and ECM from Schwann cells ^57^. In this study we tested the capability of Schwann Cells to synergize ECM production with hMSCs. Human MSCs can produce various ECM components such as fibronectin, laminin, collagen I and Tenascin-C ^58^, which can act as a substrate for Schwann cells. Co-culture of Schwann cells and hMSCs increased fibronectin deposits by 5-fold than when cultured alone. The Schwann cells were also able to add greater laminin components to the ECM in co-culture than when the hMSCs are cultured alone. In addition, the ECM fibrils aligned with the direction of the TiSAMP patterning. Certain ECM proteins are well known to assist nerve regeneration. During development, the expression of laminin and fibronectin coincides with migration of neurons ^59^. Laminin can guide axons and neurites, while fibronectin has a role in regulating cell differentiation, migration and adhesion ^60–63^. Various endogenous cells have been known to infiltrate scaffolds ^30,46,64,65^ and may contribute to their ECM composition. Since the TiSAMP chemical processing aided depositing of ECM from cultured cells, infiltrating endogenous cell types may deposit ECM which could aid regeneration. Following the trend of axon regrowth, all the implanted scaffold sheets had equal depositing of laminin and fibronectin. Since only a small proportion of the total cells were immune cells and on average the laminin and fibronectin area were greater than the scar area, this ECM depositing by endogenous cells could have been beneficial to axon regrowth. This finding also provides parameters for investigation when implanting exogenous ECMs, since endogenous cell infiltration can contribute to the overall composition. Therefore, strategies influencing endogenous cell expression of inhibitory products in combination with biomaterials remains important.

Overall, this study demonstrates that OPF scaffolds fabricated as ridged sheets with chemical patterning can promote alignment and outgrowth of cells in a bidirectional manner. This patterning also influences the directionality of produced factors such as the ECM. When rolled and implanted into rats with spinal cord transection, the rolled scaffold sheets promote greater axon regeneration than multichannel scaffolds. There is no additional immune cell infiltration or scarring noted. ECM depositing on the scaffold by endogenous cells may contribute to the axon outgrowth. This study provides a great Regenerative Medicine tool that can be used in vitro for cell characterization or in vivo for axon guidance, neuroprosthetics and devices production, or future delivery of cells and patterned ECM.

## Acknowledgements

This publication was supported by Grants from the New Jersey Commission on Spinal Cord Research CSCR15IRG002, the National Center for Advancing Translational Sciences (NCATS) UL1 TR002377 and TL1 TR002380, Mayo Clinic Benefactor Funded Career Development Award – Regenerative Medicine Initiative, and National Institute of Arthritis and Musculoskeletal and Skin Disease (R01AR07323). It was also supported by grants from the Bowen Foundation, the Kipnis Foundation, Nemitz Foundation, and the Mayo Clinic Center for Regenerative Medicine, and the New Ideas in the Natural Sciences Award from the Princeton Dean of Research.

## Data Availability

All data is available on reasonable request.

## Code Availability

Any applicable codes can be provided on reasonable request.

## Author Contributions

Conceptualization, AMS, JES, JS, AJW, NNM; performed experiments, AMS, FT, RS, SP, GW, TS, BKC, JN, JS, KL; analyzed data, AMS, FT, RS, GW; writing—original draft preparation, AMS; writing—review and editing, AMS, JES, JS, AJW, NNM (all authors had final input); visualization, A.M.S.; supervision, AMS, JES, JS, AJW, NNM.; project administration, JES, JS, AJW, NNM.; funding acquisition, JES, JS, AJW, NNM

All authors have read and agreed to the published version of the manuscript.

## Supplementary Information

### Overview of TiSAMP patterning of OPF scaffolds and implantation

**Supplementary Figure 1:**
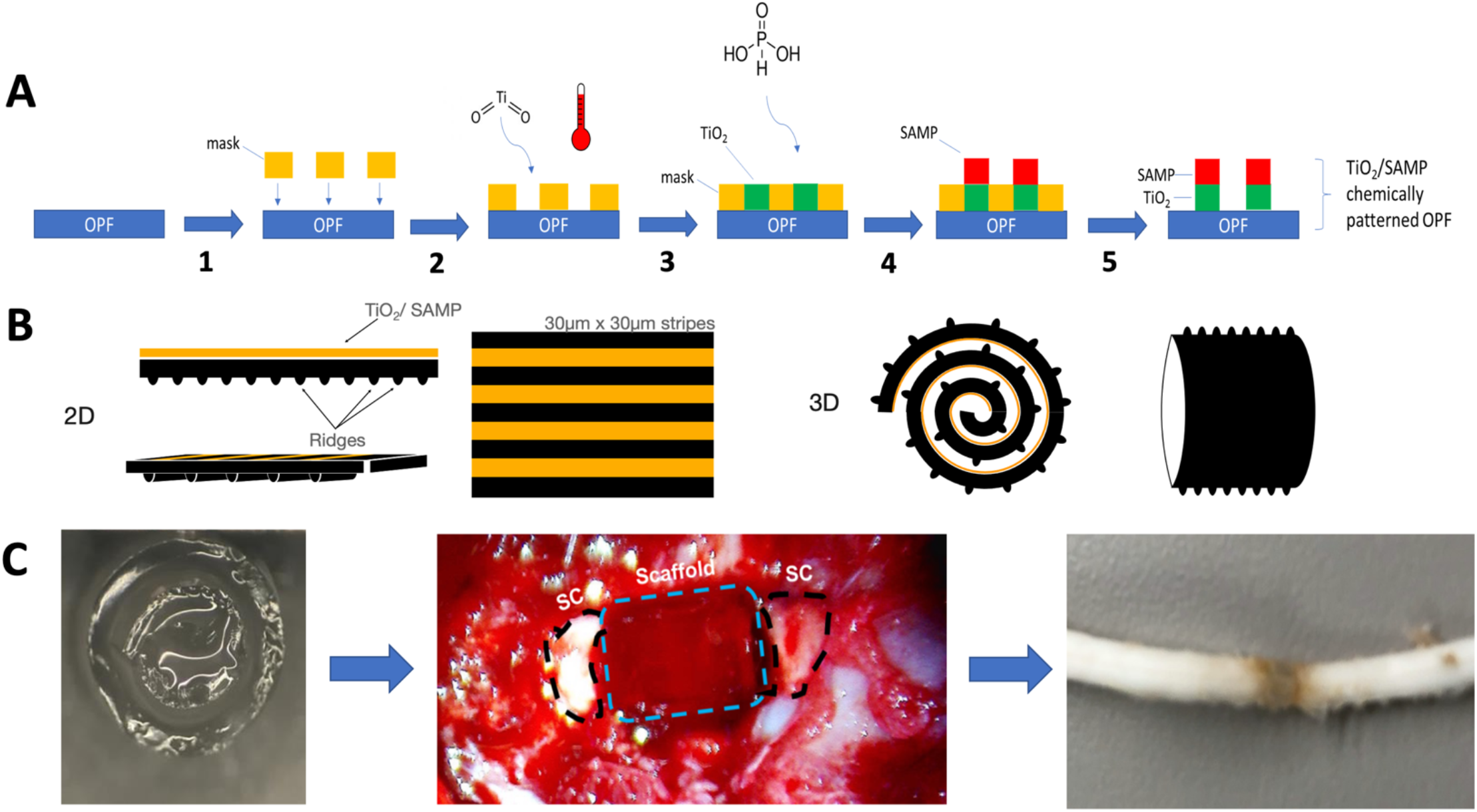
Summary of the methodology of chemical patterning on OPF scaffolds and implantation of the rolled scaffolds. (A) Schematic for debossing OPF with the micron-scale chemical pattern. The shadow mask was placed onto the OPF surface (1). The substrate was exposed to titanium dioxide and heat (2), forming a cross-linked TiO_2_ base layer. Overnight exposure to phosphonic acid (3) yielded the self-assembled layer of phosphonate (4). The mask was removed to generate 30 ⌠m x 30 ⌠m TiO_2_/SAMP pattern on the polymer surface (5). (B) The resulting OPF sheet consisted of a ridged and chemically patterned surface. The sheet could then we rolled with the ridges also acting as a spacer. (C) Once rolled, the sheet was placed in a single channel OPF scaffold to hold the structure together from implantation into the transected thoracic level 9 spinal cord. The spinal cord when removed shows integration with the spinal cord tissue.

#### Machine learning and neural networks for quantification of immune cell populations and fibrotic scar area

Histological and immunohistological analysis of tissue samples is a staple in neuroscience and regenerative medicine. After therapeutic intervention, such as after implantation of biomaterials, many aspects such as axon regeneration, immune infiltration, scarring, infiltration of endogenous cells, vascularization, and tissue preservation need to be measured ^11,31–33,35^. The in-depth analysis of these parameters using manual methods is time consuming. In the area of pathology and diagnostics, there has been a rise of digital pathology to use artificial intelligence and machine learning to inform clinical research and clinical decision making ^66,67^. Many new proprietary software have become available to establish pathology workflows informed by machine learning, however the uptake in basic science and pre-clinical research has been slower. In addition, these proprietary software can be costly for many research laboratories budgets ^68^. In this paper we demonstrate how a free open-source software, QuPath, can be used to develop machine learning algorithms to analyze immunostained and histologically stained samples to determine level of immune cell infiltration and fibrotic scarring after scaffold implantation.

QuPath has 3 types of supervised machine learning algorithms built in: random forest classifiers, K-nearest neighbor classifier, and neural networks. Random forest classifiers are computationally less extensive and typically tabular results can be produced without extensive configuration of the classifier by the user ^69,70^. Random forest classifiers depend on network of decision trees that are together are used to determine an outcome. This process has the disadvantage that it can overfit the data. K-nearest neighbor classifiers are one of the earliest algorithms used in medical research for pattern and object recognition ^70^. K-nearest neighbor uses nonparametric clustering where the classification is based on the number of k neighbors. The disadvantage of K-nearest neighbor is that processing time becomes longer for larger number of classification and data, therefore having limited scalability. Neural networks are useful when processing large number of features and allows for more complicated classification tasks ^69^. Neural networks try to mimic the brain, where a network of artificial neurons use inputs to assign mathematical weights to predict outcomes. If not monitored, like other approaches, neural networks can overfit data. However, it has been argued that using neural networks for image processing to make clinical decisions was more accurate and required less input than random forest classifiers ^71^.

Quantification of histology can be time consuming and acts as a barrier for production of in-depth quantitively studies of regenerative medicine approaches in a timely manner. In this study, we used neural networks to quantify immune cell infiltration (CD45+ and Iba-1+ cells) and fibrotic scar area (using Trichrome staining) through quarter lengths of implanted scaffolds. Once trained, the application of the learned algorithm allows for fast, non-bias quantification of large image sets.

### Neural Networks for quantification of immune cell infiltration into implanted scaffolds form micrographs of immunostained fluorescence images

For the quantitative analysis of immune cell infiltration in each scaffold, we implemented a neural network inside the open-source image analysis software QuPath. First, we created a new QuPath project by loading all the immunostained fluorescence images and setting their image type to “Fluorescent” inside QuPath to facilitate the following differentiation of the different channels. The channel names were set to DAPI, Iba1 and CD45. Next, we created a pixel classification thresholder which outlines the borders of the tissue sample per slide based on the DAPI channel pixel intensities. This allowed us to create accurate regions-of-interest for the later classifiers. We set the minimum object size to 1000 μm and the minimum hole size to 15000 μm to produce the most accurate results when measuring the area. To now obtain cell counts for the tissue regions, we implemented a cell detection watershed classifier. This classifier created outlines for each nuclei, using again the pixel intensities of the DAPI channel with a minimum area of 10 μm, maximum area of 150 μm and a pixel intensity threshold of 35.

As the performance of machine learning classifiers can be increased by careful selection of fitting features, we calculated additional features for each cell to obtain more accurate results. We considered two classes of features: shape and intensity. Shape features included area, length, circularity and solidity, while intensity features included the mean, minimum, maximum, standard deviation and median values of pixel intensities of the CD45 and Iba-1 channel of the previously detected cells.

To create a training set for the machine learning classifier, an investigator manually labeled cells on randomly chosen images as positive for CD45, positive for Iba-1, positive for both or negative. We then implemented two neural networks inside of QuPath, one for the detection and classification of CD45+ cells and one for Iba-1+ cells. The neural networks had the same architecture with 3 layers and 12 neurons in the hidden layer. We trained them for 1000 iterations and the RPROP (resilient backpropagation) optimization algorithm for achieve increased performance on the gradient descent.

After training we combined both neural networks to create a composite classifier which is capable to differentiate between cells positive for CD45, Iba-1, both antibodies or negative for both.

To further automate the whole analysis process, we combined all the steps described above in a computer script written in the scripting language groovy. Executing this script directly inside QuPaths build-in Script Editor allowed us to pipeline the whole process and further reduce the chance of introducing bias into the process.

Lastly, all measurements were exported as a csv file for statistical analysis.

**Supplementary Figure 2:**
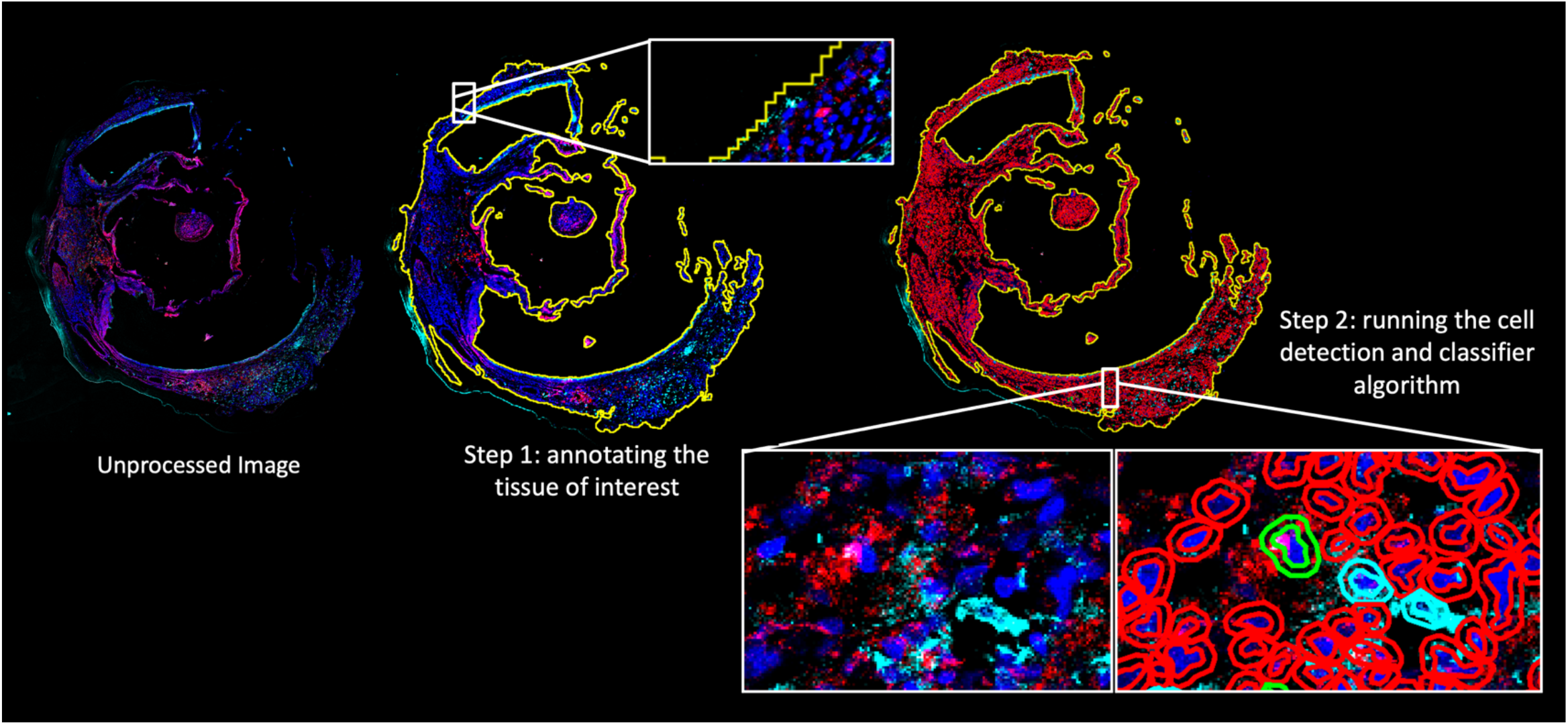
Machine learning identification of immune cell infiltration into the scaffold following implantation. Tissue sections containing implanted scaffolds were stained at quarter lengths with CD45 (cyan) and Iba-1 (red) antibodies. First the tissue of interest was annotated using a threshold algorithm to identify the borders of the tissue of interest. Training examples were created using single cell annotations as well as region wide annotations combined with a filter to only include objects identified as cells by the previous cell detection step. The number of training cells was approximately 20,000. This cell detection and classifier algorithm was ran on the whole data set to identify negative, positive, double positive labelled cells. Measurements for tissue area, total cell count, number of CD45+ cells, Iba-1 + cells, and cells positive for both antibodies was then exported for statistical analysis in Prism GraphPad Software.

### Color deconvolution based fibrotic scar differentiation from whole slide micrographs of brightfield images

For the analysis of the trichrome stained brightfield images we created a new QuPath project, again loading the images and setting their image type to “Brightfield” this time inside QuPath. We first performed a color deconvolution step by estimating the stain vectors of the images. For this step, representative regions of the images were selected which included stained tissue as well as background. This allowed to separate the red and blue stains more precise to differentiate the tissue later. Next, we created an outline of the tissue based on a pixel threshold classifier, similar to the procedure for the immunofluorescent images. The channel for the pixel threshold classifier was set to be the average pixel values over all the channels.

For the classification of scarred tissue, we created a second pixel threshold classifier to only detect fibrotic and scarred area based on the Aniline Blue stain. This stain labels collagen, which is expressed in excessive amounts in fibrotic tissue. Therefore, the intensity of the Aniline Blue stain correlates with the density of collagen in the tissue. The threshold for scar detection was internally validated by comparing the results with other images of trichrome-stained tissue and empirically set to 0.8.

The two classifiers in combination were used to first label the tissue of interest and then classify the scarred portion of the tissue. All the steps were again combined into a computer script which allows faster execution and reproducible results.

Final measurements were exported and percentages for fibrotic and scarred area were obtained by dividing scarred tissue area by whole tissue area.

**Supplementary Figure 3:**
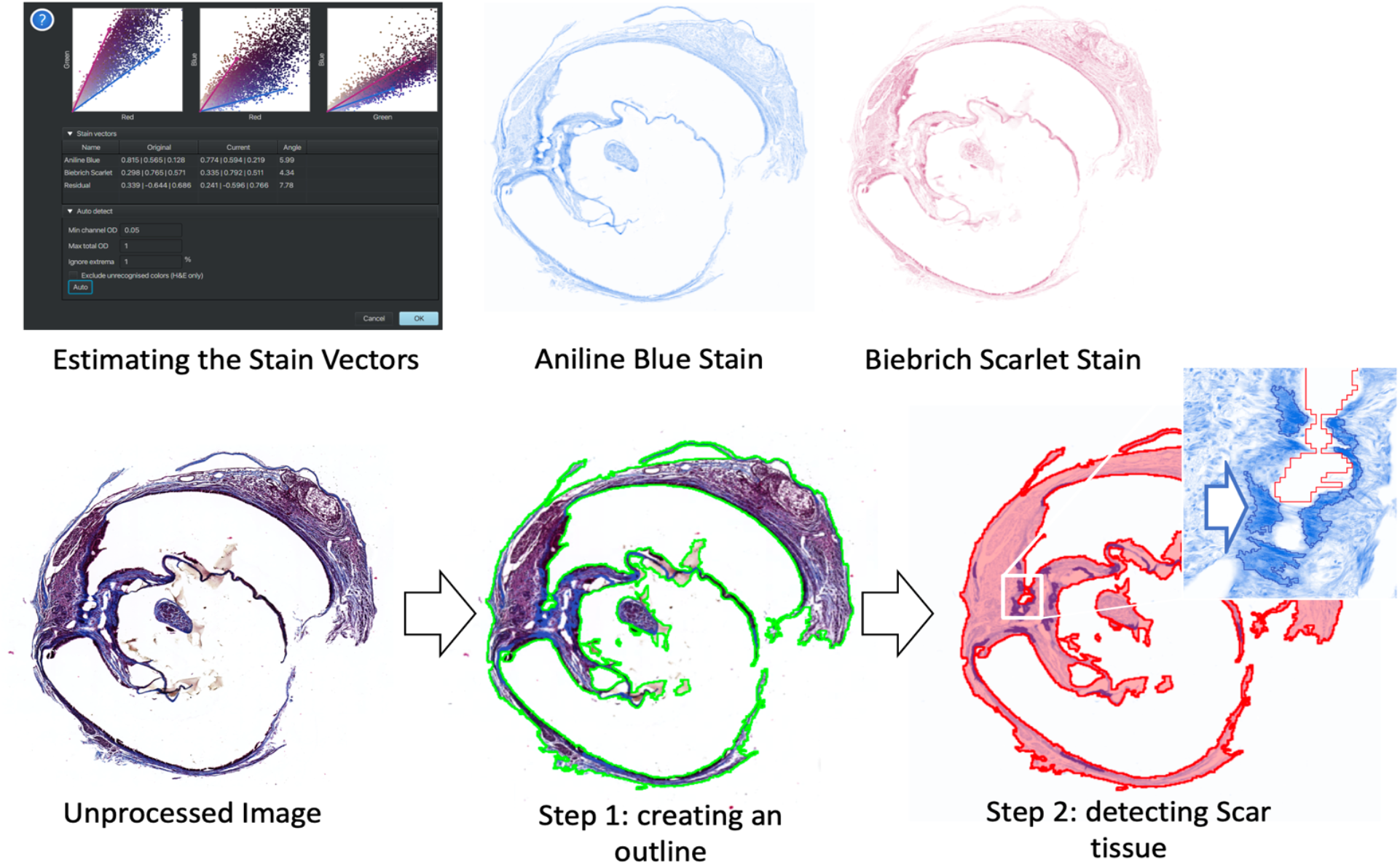
Color deconvolution based analysis of fibrotic scar area from Trichrome stained samples. Scaffold tissue samples at quarter lengths through the scaffold were stained with Trichrome histological staining method. Brightfeild scans of whole slides were taken and saved as individual files. To accurately distinguish the Aniline Blue stain (blue) from the Biebrich Scarlet stain (red), the stain vectors were estimated using pixel value representations. After this color deconvolution, a pixel threshold classifier was used to identify the boundaries of the tissue. The scar tissue was trained on a second pixel classifier designed to identify the excessive depositing of collagen stained by the Aniline Blue which is identified by the darker blue staining (arrows in step 2) than the ubiquitous collagen present in the tissue. The measurements for the whole tissue area and scar area were exported and percentages were calculated by dividing the scar area by the total area. Statistical analysis was performed in Prism GraphPad software.

